# Composition and RNA binding specificity of metazoan RNase MRP

**DOI:** 10.1101/2025.02.21.639568

**Authors:** Yuan Liu, Shiyang He, Anzie Pyo, Shanshan Zheng, Meijuan Chen, Sihem Cheloufi, Nikolai Slavov, William F Marzluff, Jernej Murn

## Abstract

Ribonuclease (RNase) MRP is a conserved RNA-based enzyme that is essential for maturation of ribosomal RNA (rRNA) in eukaryotes. However, the composition and RNA substrate specificity of this multisubunit ribonucleoprotein complex in higher eukaryotes remain a mystery. Here, we identify NEPRO and C18ORF21 as constitutive subunits of metazoan RNase MRP. Both proteins are specific to RNase MRP and are the only ones distinguishing this enzyme from the closely related RNase P, which selectively cleaves transfer RNA-like substrates. We find that NEPRO and C18ORF21 each form a complex with all other subunits of RNase MRP, stabilize its catalytic RNA, and are required for rRNA maturation and cell proliferation. We harness our discovery to identify a full suite of *in vivo* RNA targets of each enzyme, including positions of potential cleavage sites at nucleotide resolution. These findings resolve the general composition of metazoan RNase MRP, illuminate its RNA binding specificity, and provide valuable assets for functional exploration of this essential eukaryotic enzyme.

## Introduction

The maturation of 18S, 5.8S, and 28S rRNA begins with endonucleolytic cleavages of the polycistronic precursor rRNA (pre-rRNA) and continues with exonucleolytic trimming to produce the mature rRNAs essential for the formation of functional ribosomes^1^. Cleavage that splits precursor ribosomal RNA (pre-rRNA) into parts destined for incorporation into the small or the large subunit of the human ribosome is mediated by a poorly understood, evolutionarily ancient RNA-based enzyme, RNase MRP (Figure 1A)^1, 2^. This essential eukaryotic ribonucleoprotein (RNP) is known to consist of at least nine protein subunits (POP1, POP5, RPP14, RPP20, RPP25, RPP29, RPP30, RPP38, RPP40) and a catalytic RNA, termed *RMRP1*^3,4^. Like its composition, the *in vivo* function of RNase MRP is not understood well. However, its irreplaceable role in human cells is highlighted by mutations in *RMRP1* that cause various pleiotropic diseases, including cartilage-hair hypoplasia (CHH) characterized by skeletal abnormalities, immune system deficiency, and increased cancer risk^5, 6^.

**Figure 1.**
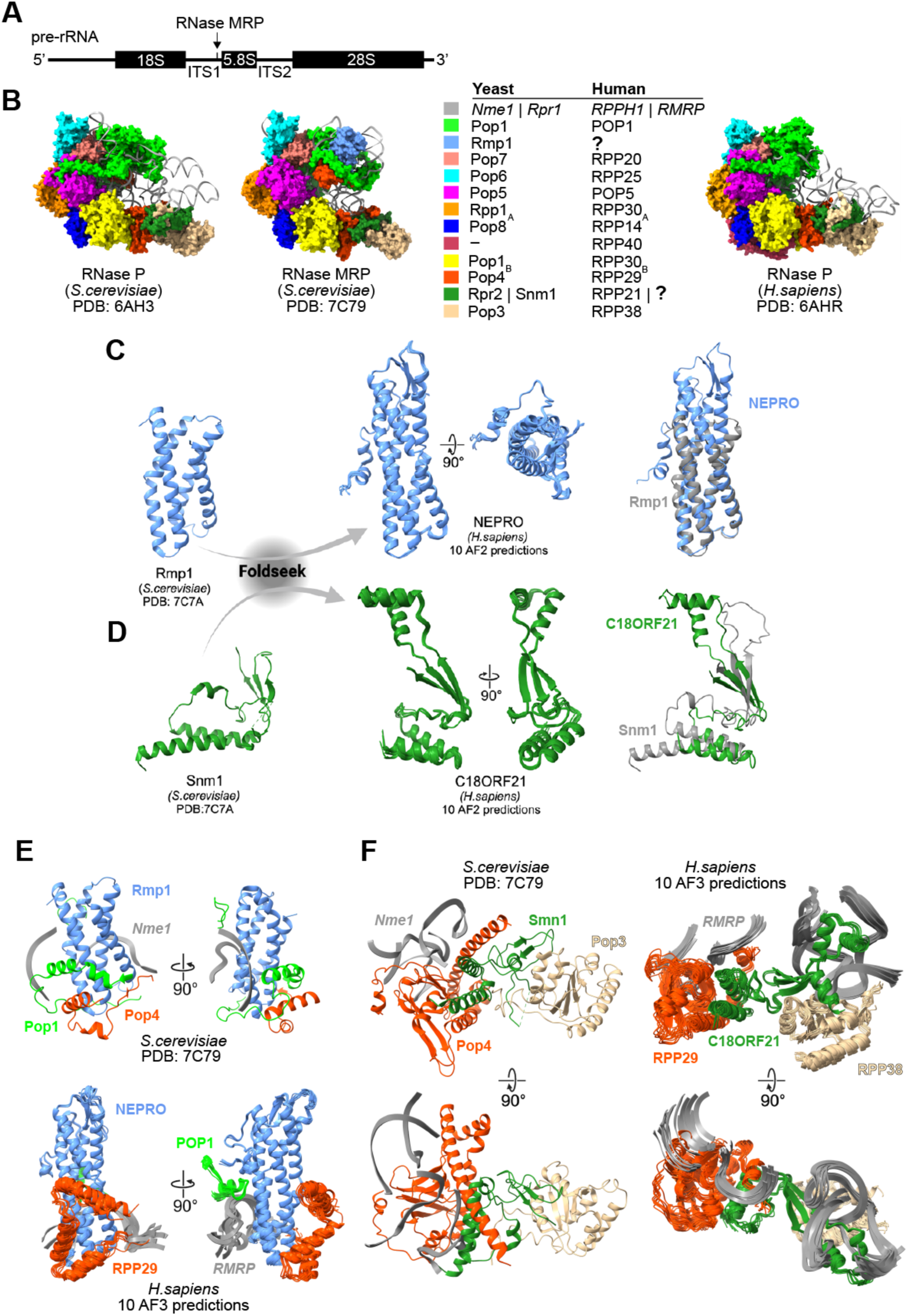
Structural homology identifies two novel candidate subunits of metazoan RNase MRP. **(A)** Schematic indicating the location of the annotated RNase MRP cleavage site within ITS1 of 47S pre-rRNA. **(B)** Cryo-EM structures of yeast RNase P (PDB: 6AH3, left), yeast RNase MRP (PDB: 7C79, middle), and human RNase P (PDB: 6AHR, right) along with listing of known RNase P and RNase MRP subunits in yeast and human. **(C, D)** Foldseek identified human NEPRO and C18ORF21 as structural homologs of yeast Rmp1 and Snm1, respectively. Cryo-EM structures of Rmp1 (C) and Snm1 (D, both PDB: 7C7A) were used as queries. Superpositions of ten AlphaFold2 (AF2) predictions of the converging segments of human NEPRO and C18ORF21 are shown. The same segments of each protein (one AF2 prediction) were also used for their alignments with the cryo-EM structures of Rmp1 and Snm1, as shown. **(E)** Interfaces of yeast Rmp1 interacting with Pop1, Pop4, and *Nme1* (PDB: 7C79; top) compared to ten AlphaFold3 (AF3) predictions of human NEPRO interacting with POP1, RPP29, and *RMRP* (bottom). Ten AF3 predictions of human RNase MRP were generated using full-length sequences of all human subunits listed in panel (B), NEPRO, and C18ORF21. The predictions were aligned on NEPRO. Only the converged segments of NEPRO and its interacting subunits are shown. **(F)** As in (E) but showing interfaces of yeast Snm1 interacting with Pop4, Pop3, and *Nme1* (PDB: 7C79; left) compared to ten AF3 predictions of human C18ORF21 interacting with POP1, RPP29, and *RMRP* (right). The predictions of the human RNase MRP were aligned on C18ORF21 and only the converged segments of C18ORF21 and its interacting subunits are shown.

RNase MRP is structurally and evolutionarily related to another essential RNA-based enzyme, RNase P, which is required for the maturation of the 5’ end of precursor transfer RNA (pre-tRNA) in all three domains of life^4, 7^. Structural studies in yeast reveal that both RNPs use a similar catalytic core within the RNA moiety and share most protein subunits^4, 8, 9^. However, several RNA elements and a handful of protein subunits distinguish RNase MRP from RNase P in yeast^8, 9^. These differences are thought to underlie the enzymes’ distinct modes of RNA substrate recognition: ’measuring’ the L-shaped pre-tRNA-like structures by RNase P and recognition of a short consensus sequence in single-stranded RNA substrates by yeast RNase MRP^8, 10–13^. Whereas one human RNase P-specific protein has been identified (RPP21; orthologous to Rpr2 in yeast)^14^, no human RNase MRP-specific proteins were found, although two such proteins, Rmp1 and Snm1, exist in yeast^4, 15, 16^. The apparent absence of RNase MRP-specific subunits has puzzled the field for several decades, leading to the current lack of even rudimentary understanding of this intriguing enzyme in higher eukaryotes.

A vexing question is about the sheer existence of RNase MRP in higher eukaryotes: why would a simple hydrolytic cleavage of one RNA substrate require a half-megadalton RNA-based catalyst in human cells? RNase MRP is thought to have emerged in eukaryotes as an RNase P-like enzyme to fulfill a new requirement for pre-rRNA processing^7^. This could justify its dedication to processing of pre-rRNA in the earliest eukaryotic organisms about two billion years ago^17^. However, it is difficult to envision that RNase MRP would sustain its single-substrate specificity in higher eukaryotes, especially considering that at least two additional RNase MRP substrates, *CLB2* and *CTS1* mRNAs, have been identified in yeast and that similar site-specific cleavages of pre-rRNA are catalyzed by much smaller protein-only complexes in mammalian cells^18–20^. The persistence of RNase MRP conceivably reflects a need for processing of multiple RNA substrates, likely with some level of sequence specificity^10^. Defining these and the purpose of their processing by RNase MRP should help decipher the *in vivo* function of this RNP.

Here we use structural predictions to discover two RNase MRP-specific protein subunits in metazoans. We validate that NEPRO and C18ORF21 are unique subunits of mammalian RNase MRP via analyses of their protein–protein and protein–RNA interactions, critical roles in pre-rRNA processing, and requirement for cell proliferation. We leverage our discovery to distinguish RNase P from RNase MRP and define complete repertoires of their *in vivo* RNA targets along with positions of putative cleavage sites at nucleotide resolution. These results shed light on the composition and RNA binding specificity of metazoan RNase MRP.

## Results

### Structural homology identifies two novel candidate subunits of metazoan RNase MRP

Recent cryogenic-electron microscopy (cryo-EM) studies in *S.cerevisiae* have revealed a considerable overall architectural similarity between RNase P and RNase MRP RNPs, as well as the preserved catalytic RNA core inherited from their common ancestor (Fig. 1B)^8, 9^. In addition, these analyses revealed incorporation of new RNA elements as well as two protein subunits, Rmp1 and Snm1, that contributed to the establishment of yeast RNase MRP as an enzyme with substrate specificity distinct from RNase P. It is unclear whether putative orthologs of the yeast RNase MRP-specific proteins might be components of metazoan RNase MRP.

As sequence-based searches for human orthologs of Rmp1 or Snm1 found no obvious candidates, we asked whether such proteins might exist based on their structural homology. We used Foldseek to query the cryo-EM structures of the two yeast proteins within RNase MRP against thousands of experimentally determined or computationally predicted protein structures^21^. This search identified the N-terminal helical bundle of the little studied nucleolar protein NEPRO^22–24^ as distinctly the closest structural match to the yeast Rmp1 in all inspected metazoans species, despite the limited amino acid sequence conservation (Figures 1C and S1A, B). A similar search of the yeast Snm1 protein revealed the most extensive homology with two human proteins: the RNase P-specific subunit RPP21 and an uncharacterized protein C18ORF21 (Figure 1D). Interestingly, in yeast, the RNase MRP-specific Snm1 is structurally most homologous with the RNase P-specific Rpr2, and both proteins are similarly nestled in their RNP complexes between Pop4 and Pop3 (Figures 1B and S1C)^8, 9^. It is thus plausible that RPP21 and its homologous C18ORF21 occupy the same relative positions in metazoan RNase P and RNase MRP, respectively (Figure S1D-S1F).

We used AlphaFold to generate structure predictions of human NEPRO and C18ORF21 in complex with the previously annotated subunits of the human RNase MRP (Figure S2A)^3^. These predictions suggested that twelve protein subunits, including NEPRO and C18ORF21, sequentially contact one another and tightly wrap around *RMRP*, much like the orthologous proteins of the yeast RNase MRP interlink together to stabilize *Nme1* (Figures 1B and S2A)^8, 9^. However, unlike the hook-shaped architecture of RNase MRP in yeast, the predictions of human RNase MRP uniformly indicate a more closed, rigid conformation of the RNP that is stabilized by extensive interactions of NEPRO and C18ORF21 with *RMRP* and with one another (Figure S2A-C). Additionally, all predictions show that the structures of the catalytic (C) domain in the human *RMRP* and yeast *Nme1* are nearly identical, while suggesting entirely different topologies of the specificity (S) domain between the two species (Figure S2D)^3, 8, 9, 25^.

A comparison of the cryo-EM structure of the *S. cerevisiae* RNase MRP holoenzyme with structural predictions of its human counterpart point to striking similarities in the way that either RNase MRP-specific subunit is integrated into the enzyme. In both human and yeast, the extended N-terminal domain of RPP29/Pop4 wraps around one side of the helical bundle of NEPRO/Rmp1 while POP1/Pop1 contacts the bundle from the opposite side (Figures 1E and S2A). All generated predictions further suggest intimate interactions of NEPRO with *RMRP* and position NEPRO atop the catalytic RNA pseudoknot (Figures 1E and S2C). This resembles the placement of Rmp1 in yeast RNase MRP where it forms part of the substrate-binding groove and facilitates recognition of single-stranded RNA substrates (Figure S2B)^8, 9^. At the other end of the enzyme, Snm1 interacts with Pop4 and Pop3 in yeast in a similar fashion as C18ORF21 is predicted to interact with RPP29 and RPP38 in humans (Figures 1B, F and S2A). However, compared to Snm1, the predictions suggest considerably more extensive interaction of C18ORF21 with the RNase MRP RNA (Figures 1F and S2B, C).

Together, these analyses identify two novel candidate subunits of the metazoan RNase MRP and predict their RNP-specific structural roles.

### NEPRO and C18ORF21 are constitutive components of RNase MRP but not RNase P

To confirm the predicted subunit status of NEPRO and C18ORF21 in RNase MRP, we sought to validate the association of each protein with the known subunits of the enzyme in human and mouse cells. We used affinity-purification mass spectrometry (AP-MS) to enable enrichment and detection of proteins that stably associate with different ectopically expressed tagged RNase P and/or RNase MRP subunits, which we used as baits. Tandem AP-MS analysis of the complex formed by RPP25 in HEK293T or mouse embryonic stem cells (mESCs) found among the most highly and specifically enriched proteins all subunits annotated as common to both enzymes, the RNase P-specific RPP21, as well as NEPRO and C18ORF21 (Figure 2A, B). These proteins were also overrepresented in other complexes that we purified from mESCs, except that Rpp21 was not detected in the Nepro- or C18orf21-nucleated complexes and, vice versa, Nepro or C18orf21 were not detected in the Rpp21 complex (Figures 2C-E). We used co-IP/western analysis to validate the coexistence of Nepro and C18orf21, but the absence of Rpp21 within the same protein complex (Figure 2F).

**Figure 2.**
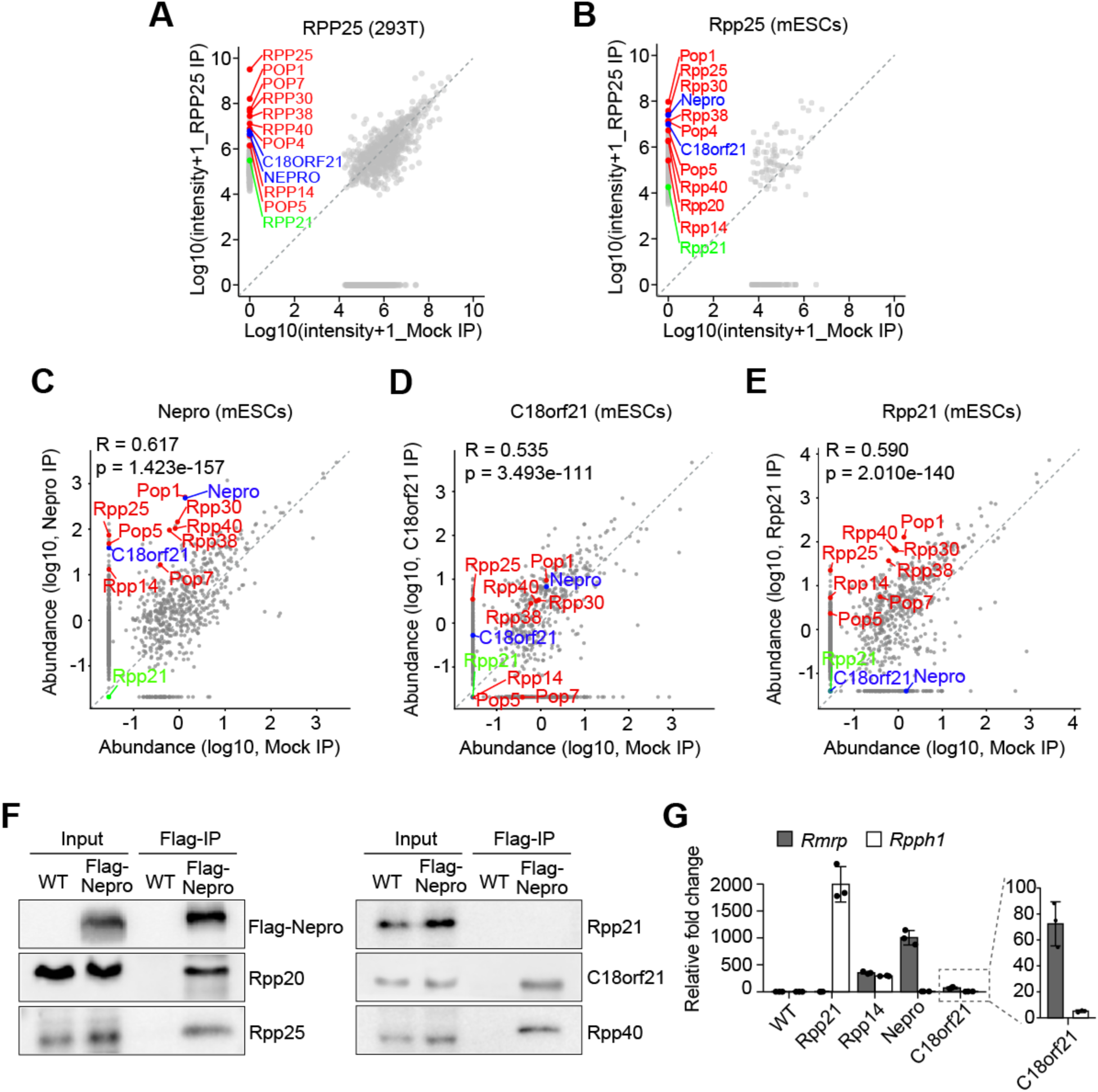
NEPRO and C18ORF21 are constitutive components of RNase MRP but not RNase P. **(A, B)** Results of mass spectrometry analyses comparing summarized intensities for each protein detected in tandem affinity-purified protein complex (anti-Flag IP followed by anti-HA IP) of RPP25 and mock control in HEK293T cells (A) and mESCs (B). RNase MRP protein subunits in common to RNase P are highlighted in red, the RNase P-specific RPP21 is in green, and the two newly identified RNase MRP-specific subunits, NEPRO and C18ORF21, are in blue. **(C-E)** Results of mass spectrometry analyses comparing abundances of each protein detected in affinity-purified protein complex (anti-Flag IP) of Nepro (C), C18orf21 (D), or Rpp21 (E) and mock control in mESCs. Protein subunits of RNases P and MRP were highlighted as in (A, B). **(F)** Co-IP of endogenous RNase P and/or MRP protein subunits with Nepro from lysates of mESCs stably expressing 3xFlag-Nepro. Precipitated proteins were detected by western blot analysis. **(G)** RNA immunoprecipitation (RIP) analysis of *Rmrp* and *Rpph1* co-precipitated with Flag-tagged Rpp21, Rpp14, Nepro, or C18orf21 from lysates of mESCs. Relative fold change of *Rmrp* or *Rpph1* in each RIP experiment compared to wild-type (WT) control was determined using qPCR (n = 3). Data are shown as mean ± SD.

Given that RNases P and MRP use distinct catalytic RNA molecules, *RPPH1* and *RMRP*, respectively, we reasoned that any RNase MRP-specific protein subunit should preferentially associate with the latter. Markedly, RNA immunoprecipitation (RIP) analyses of Nepro or C18orf21 in mESCs revealed an overwhelming binding preference for *Rmrp* compared to *Rpph1* by either protein, whereas Rpp21 showed the expected specificity for *Rpph1* (Figure 2G). Thus, the strong associations of both Nepro and C18orf21 with all known RNase MRP subunits, as well as with one another and with the RNase MRP RNA, support our structural analyses and identify Nepro and C18orf21 as specific protein subunits of metazoan RNase MRP.

### The *in vivo* RNA-binding specificity of RNase MRP

We asked whether the specific association of the newly identified subunits with RNase MRP but not RNase P could be leveraged to shed light on the enigmatic RNA-targeting specificity of RNase MRP in mammalian cells. We considered that in yeast, a short single-stranded segment of the pre-rRNA’s ITS1 region - a known substrate of the yeast RNase MRP - is accommodated deep within the catalytic center of the enzyme via direct contacts with several protein subunits, in addition to interactions with the *Nme1* RNA (Figures 1A and S2B)^8^. In fact, an earlier study of the structural organization of yeast RNase MRP demonstrated that many interactions between its protein subunits and substrate RNA could be successfully captured *in vitro* using UV light-mediated crosslinking^13^. To detect similar protein-RNA substrate interactions of mammalian RNase MRP *in vivo*, we carried out individual nucleotide resolution UV crosslinking and immunoprecipitation (iCLIP)^26, 27^ of tagged RNase P and/or MRP subunits in mESCs.

We achieved co-purification of most RNP protein subunits crosslinked to the catalytic RNA as well as RNA substrates by using stringent but nondenaturing conditions, which were previously shown to preserve relatively stable snRNP complexes (Figure S3A)^28^. High concentration of salt and detergents minimized co-purification of weakly associated proteins but preserved the more stable inter-subunit interactions of RNases P or MRP, as evident from multiple radioactive bands whose pattern was similar between iCLIP experiments (Figures 3A and S3A)^27, 28^. We prepared cDNA libraries from a broad distribution of crosslinked RNP complexes to maximize the diversity of recovered protein-RNA interactions (Figure S3A).

**Figure 3.**
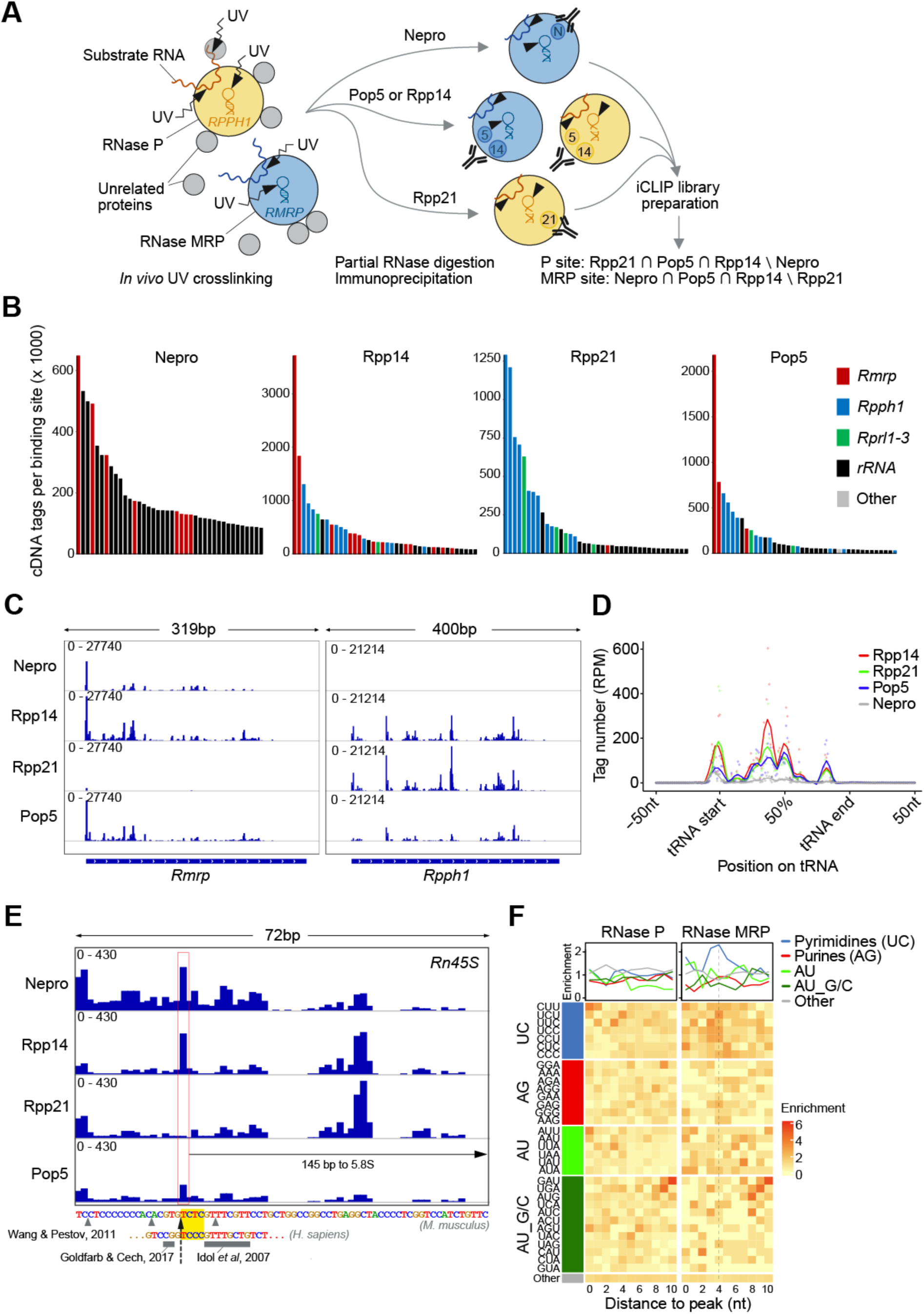
The *in vivo* RNA-binding specificity of RNases P and MRP. **(A)** Schematic representation of the iCLIP strategy to determine RNA-binding specificities of RNases P and MRP. **(B)** RNA binding sites identified by the largest number of unique cDNA tags in each dataset. Top 40 binding sites were ordered by the descending number of iCLIP tags from left to right. Gencode vM25 annotations of individual sites are shown. **(C)** Crosslinking profiles on *Rmrp* and *Rpph1* shown for each iCLIP dataset. Numbers in each trace are counts per million (CPM) values. **(D)** Proportional metatranscript analysis of iCLIP data showing the positional frequency of crosslink events on tRNAs. Data points represent normalized crosslink events summarized over every percent of the tRNA length. RPM, reads per million. **(E)** Crosslinking profile in the ITS1 segment of the mouse *Rn45S* transcript (NR_046233.2) highlighting the sole RNase MRP-specific site (red rectangle) and its distance from the 5.8S rRNA. Previously mapped RNase MRP cleavage sites are indicated separately on the mouse transcript (gray arrowheads)^37^ and the orthologous section of the human transcript (gray rectangles)^34, 36^. The yellow rectangle and the black dashed arrow indicate putative RNase MRP consensus recognition sequence and cleavage site, respectively. **(F)** Heatmaps illustrating positional frequencies of the indicated sets of trimers 10 nts downstream of the RNase P (left) and RNase MRP (right) binding site maxima. Plots above the heatmaps profile the mean enrichment of the different sets of trimers.

Data analysis revealed several thousand protein–RNA interactions, the majority of which were identified with high reproducibility in each iCLIP dataset (Figure S3B). Binding sites with the largest numbers of unique crosslink events were uniformly located on the catalytic RNAs of RNases P and/or MRP, with crosslinks from Nepro iCLIP mapping selectively to *Rmrp* and those from Rpp21 iCLIP mapping to *Rpph1*, whereas the analysis of Rpp14 or Pop5 showed strong binding to both catalytic RNAs (Figure 3B). A further inspection of the crosslinking to *Rmrp* or *Rpph1* revealed complex but very similar RNA binding patterns across the different iCLIP datasets, consistent with the capture of RNP-wide and not merely subunit-specific protein-RNA interactions (Figures 3C and S3A). For a crosslinking peak to be ascribed to either RNase P or RNase MRP, we required that it be called in 1) exactly three out of four independent iCLIP datasets and 2) one but not both enzyme-specific subunit datasets, i.e., Rpp21 or Nepro but not both (Figure 3A). This requirement secured validity of binding sites by disregarding those that were either non-specific (e.g., called in each iCLIP dataset due to adventitious binding of a highly expressed transcript) or were specific to the subunit but not the RNP complex (e.g., Nepro possibly occurs in cells outside of RNase MRP and may bind RNA independently)^22^.

In addition to binding of the catalytic RNA, we detected interactions of RNases P and MRP with several other RNA species, including rRNA, tRNA, mRNA, as well as with different types of short and long non-coding RNA (Figures 3B and S3C, D). To assess the extent to which these interactions may indicate recognition of specific RNA substrates by RNases P and MRP, we first examined the binding of each enzyme to their known RNA targets. We found that Rpp14, Rpp21, Pop5 but not Nepro iCLIP cDNAs mapped extensively to multiple positions along the length of tRNAs, in line with pre-tRNA being a dominant substrate of RNase P but not RNase MRP (Figures 3D and S3E)^3, 7, 29^. This included a prominent crosslinking peak just upstream of the annotated mature tRNA segments, which agrees with structural studies of RNase P in different organisms that show or predict critical protein interactions with the 5’ leader of the pre-tRNA substrates at the catalytic center^11, 12^. We also examined interactions with another documented RNA substrate of RNase P, *Malat1*, whose conserved triple helical 3’ end is produced by recognition and cleavage of a tRNA-like structure by RNase P^30, 31^. Strikingly, we found a dominant RNase P-specific binding site on the ∼7-kb *Malat1* transcript precisely at the annotated RNase P cleavage site within a short A-rich tract (Figure S3F).

While RNase P recognizes its substrates primarily based on their shape and by acting as a molecular ruler^11, 12, 32^, little is still known about substrate recognition by RNase MRP, aside from its preference for a loosely defined consensus sequence in yeast, 5’-*RCRC-3’ (where * is the cleavage site and R is a non-obligate purine), in which a cytosine at position +4 is particularly important^8, 33^. To gauge whether iCLIP might, as in the case of RNase P, identify RNase MRP substrates based on their crosslinking pattern, we inspected pre-rRNA as the only *bona fide* RNase MRP substrate in mammalian cells (Figure 1A)^34^. Despite high background signal, we identified a single RNase MRP-specific crosslinking site on the 13-kb pre-rRNA transcript based on the peak presence in Nepro, Rpp14, and Pop5 but not Rpp21 iCLIP datasets (Figure 3E). Markedly, this site was located in the 1-kb ITS1 segment, 145 nts upstream of 5.8S rRNA, in the vicinity of previous mappings of the RNase MRP cleavage site in the mouse and orthologous sequences in the human pre-rRNA (Figures 1A and 3E)^1, 34–37^.

We also noticed that the RNA sequence immediately downstream of the crosslinking site in ITS1 may fulfill the reported substrate recognition requirement of yeast RNase MRP, containing cytosines at positions +2 and +4 relative to the peak at position +1 (Figure 3E)^8, 33^. In fact, a consideration of all identified RNase P- or RNase MRP-specific binding sites indicated that the latter were distinctly enriched in U/C-containing motifs just downstream of the binding peak, showing maximum enrichment at position +4, whereas RNase P-specific peaks showed no such trend (Figure 3F). This analysis suggests that iCLIP may be, as in the case of known RNA substrates (Figures 3D, E, and S3E, F), capturing crosslinks right at the sites of cleavage in the candidate RNA substrates (Figure 3F). Future studies are warranted to investigate potential RNase P- or MRP-dependent cleavage of the identified target RNAs, along with the biological implications of such cleavage events.

### NEPRO and C18ORF21 are required for pre-rRNA processing by mammalian RNase MRP

To understand the contribution of NEPRO and C18ORF21 to the enzymatic activity of mammalian RNase MRP, we evaluated the effects of their depletion in human or mouse cells. We rapidly and nearly completely removed NEPRO via targeted protein degradation using a conditional degradation tag (dTAG) approach, while we most efficiently depleted C18ORF21 by siRNA-mediated gene silencing (Figure 4A, B)^38, 39^. Both proteins distinctly localized to nucleoli, as would be expected from subunits of the overwhelmingly nucleolar RNase MRP^40, 41^, though neither exerted any overt effect on nucleolar size or number upon depletion in HEK293T cells (Figure 4C, D). Moreover, depletion of NEPRO or C18ORF21 led to a substantial decrease in the level of *RMRP1* but not *RPPH1*, consistent with direct, stabilizing interactions that each of the two proteins are predicted to form with *RMRP1* within the RNase MRP RNP complex (Figures 1E, F, S2C, and 4E-G).

**Figure 4.**
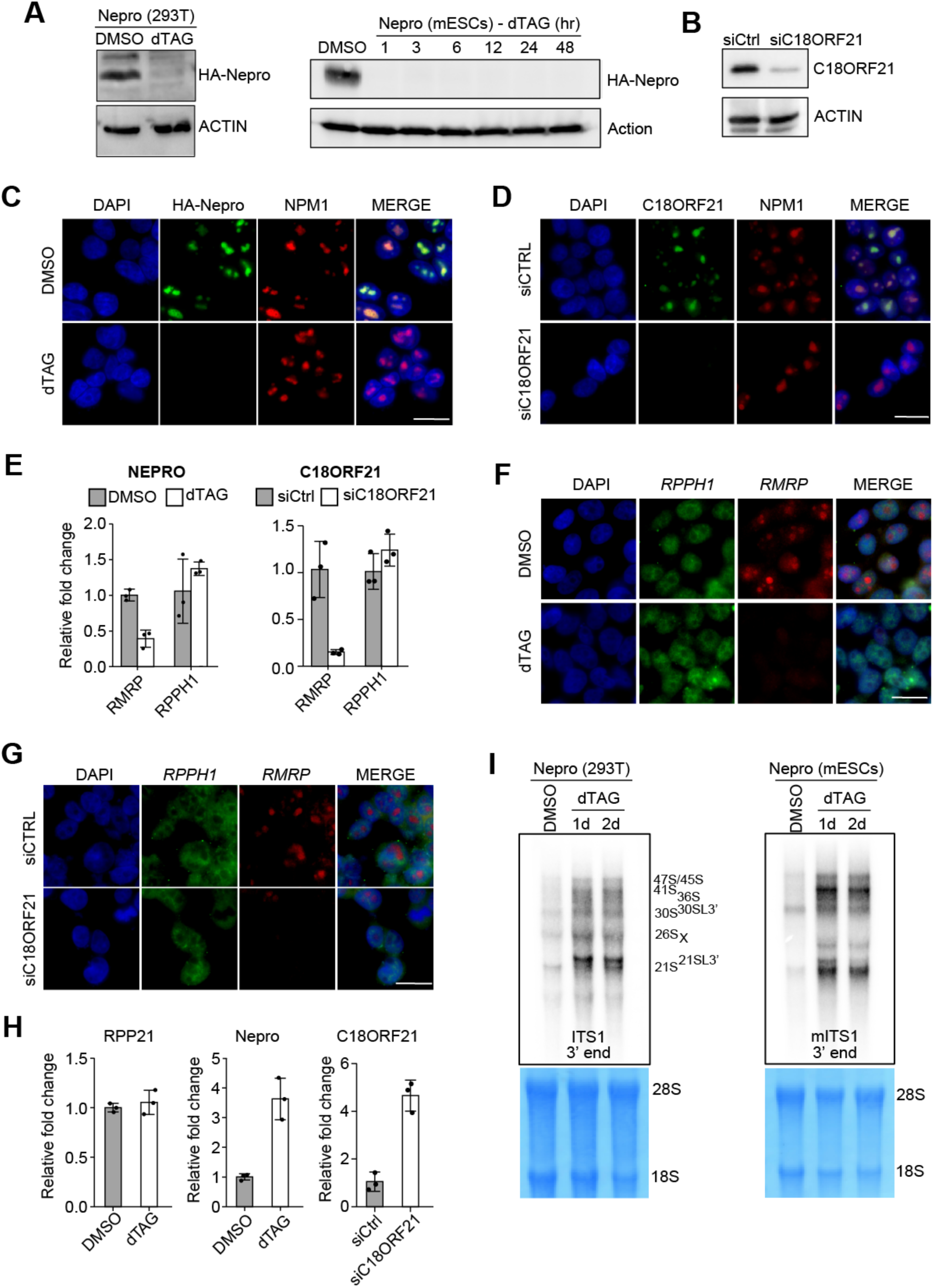
Mammalian RNase MRP requires NEPRO and C18ORF21 for pre-rRNA processing. **(A)** Depletion of HA-tagged Nepro in HEK293T cells (left) or mESCs (right) engineered for rapid depletion of the protein subunit using the dTAG system^38, 39^. HEK293T cells were treated with dTAG for 48h and mESCs as indicated prior to analysis by immunoblotting. Actin serves as a loading control (n = 3). **(B)** Depletion of C18ORF21 in HEK293T cells using siRNA at 2 days post transfection (n = 3). **(C)** The engineered HEK293T cells from (A) were analyzed by immunofluorescence using anti-HA antibody after treatment with DMSO or dTAG for 2 days. Antibodies targeting endogenous nucleophosmin (NPM1) and DAPI were used to visualize nucleoli and nuclei, respectively. **(D)** As in (C), only that C18ORF21 was depleted in wild-type HEK293T cells with siRNA for 2 days prior to the analysis. **(E-G)** Depletion of NEPRO, as in (C), or C18ORF21, as in (D), reduces steady-state levels of *RMRP* but not *RPPH1*. Cells were analyzed by qPCR (E) or by fluorescence in situ hybridization (FISH)(F, G) (n = 3). Data in (E) are shown as mean ± SD. **(H)** qPCR analysis of the RNase MRP-targeted site in pre-rRNA in cells lacking RPP21, Nepro, or C18ORF21 (n = 3). Data are shown as mean ± SD. **(I)** HEK293T or mESCs engineered for rapid depletion of Nepro, as in (A), were treated with DMSO or dTAG for 1 or 2 days, then total RNA was extracted and analyzed by northern blotting using a probe that recognizes a 3’ region of ITS1 in pre-rRNA. Methylene blue stain of 28S/18S serves as a loading control. Scale bars in (C), (D), (F), and (G) are 25 µm.

Reduction in the level of an essential catalytic RNA prompted a question of whether the processing of its key cellular substrate, pre-rRNA, might also be affected. A qPCR analysis of nuclear RNA using primers that amplify a region spanning the RNase MRP cleavage site found little difference upon depletion of the RNase P-specific RPP21 protein, but indicated severely compromised cleavage when either NEPRO or C18ORF21 were depleted (Figures 4H and S4A). Northern blot analyses confirmed a block in early pre-rRNA processing upon depletion of NEPRO in HEK293T and mES cells, showing accumulation of 47S/45S rRNA and other RNA precursors consistent with a defect in cleavage of the ITS1 segment by RNase MRP (Figure 4I)^34, 42^. Accumulation of ITS1-retaining pre-rRNA was also observed through fluorescence *in situ* hybridization analysis, which revealed a substantially increased ITS1 signal in nucleoli of NEPRO-depleted cells (Figure S4B).

Finally, we found that the rapid depletion of NEPRO or C18ORF21 caused a proliferative arrest but preserved cellular viability for several days until termination of the experiment (Figure S4C and data not shown). Though unexpected, the observed enduring cell viability agrees with previous studies demonstrating long-term survival of cells arrested by inhibition of rRNA biogenesis, including via inhibition of rRNA transcription and downstream processing^43–46^. Taken together, our structural analyses (Figure 1), studies of protein-protein and protein-RNA interactions (Figures 2 and 3), and *in vivo* functional studies (Figure 4) elucidate the composition and function of RNase MRP in metazoans, paving the way to a better understanding of this critical RNA-based regulator of ribosome biogenesis.

## Discussion

Despite its essential role in rRNA processing, ancient evolutionary origin, and association with a series of severely debilitating diseases, metazoan RNase MRP remains an enzyme of unclear composition and substrate specificity. Our discovery of two human RNase MRP-specific protein subunits, NEPRO and C18ORF21, which are conserved across metazoans, provides a critical molecular handle for structural and functional characterization of RNase MRP. This includes a solution to the long-standing problem of distinguishing the protein composition of RNase MRP from it structurally and evolutionarily closely related RNase P. To assist with these efforts, we generated cell lines for rapid, on-demand depletion of RNase P or MRP, as well as a database of candidate RNA substrates with annotated crosslinking sites of each enzyme.

We identify both human RNase MRP subunits, NEPRO and C18ORF21, as the closest predicted structural homologs of yeast Rmp1 and Snm1, respectively, despite poor sequence conservation. In fact, the similarity between the orthologous subunits is limited to the structural elements that anchor each yeast subunit into the RNase MRP complex: the N-terminal four-barrel bundle in NEPRO/Rmp1 and the N-terminal helices and beta-sheets in C18ORF21/Snm1 (Figure 1C, D). Notably, the same structural elements are consistently predicted to anchor NEPRO and C18ORF21 into the human RNase MRP complex at comparable relative positions as in the yeast complex and foster many similar interactions with the neighboring subunits and the catalytic RNA (Figures 1E, F and S2B, C). However, NEPRO, which is more than twice the size of Rmp1 in yeast, and C18ORF21 are predicted to form much more extensive contacts with the catalytic RNA than their yeast counterparts (Figures 1F and S2B, C) and directly interact with each other to ‘close’ the general hook-shaped architecture of the enzyme (Figures 1B, S2A)^8, 9^. Hypothetically, these metazoan specific protein–RNA interactions contribute to stabilization of the catalytic RNA, in line with the effects of NEPRO or C18ORF21 depletion in mammalian cells (Figure 4). The evolution of RNase MRP from its last universal common ancestor with RNase P thus appears to have continued the trajectory of becoming progressively larger and more proteinaceous in eukaryotes (Figures 1B and S2A)^8, 47^.

One pressing question regarding the biological functions of RNase MRP revolves around its RNA targeting repertoire. Despite the identification of two mRNA substrates in yeast^18, 19, 48^, the quest to understand the RNA binding specificity of RNase MRP in higher organisms has been complicated by lack of knowledge of the enzyme’s composition and its sharing of the known protein subunits with RNase P^3^. We circumvented these issues by pairing our identification of RNase MRP-specific protein subunits with iCLIP to allow for transcriptome-wide discovery of RNase P or RNase MRP-specific RNA binding sites (Figure 3A)^26, 27^. Strikingly, this analysis not only captured the known RNA substrates of RNase P (tRNAs and *Malat1*) and RNase MRP (pre-rRNA), but in every case also pointed to a prominent crosslinking peak precisely at the expected site of cleavage (Figures 3D, E and S3F). This result indicates particularly efficient *in vivo* crosslinking of protein subunits to the RNA substrate at the catalytic core of each enzyme, consistent with results of the *in vitro* crosslinking experiments with yeast RNase MRP^13^. Generalized to all identified RNase MRP target sites, such crosslinking preference would point to a pyrimidine-rich consensus sequence for mammalian RNase MRP (Figure 3E, F), hypothetically 5’-*YCYC-3’ (where * is a potential cleavage site and Y is a non-obligate pyrimidine), contrasting with the 5’-*RCRC-3’ consensus determined for the yeast RNase MRP^8, 13^.

As anticipated, we identified *Rmrp* and *Rpph1* as the most highly bound RNA molecules with over a million unique iCLIP cDNAs mapped at some of the strongest bound sites (Figure 3B). Although such read depth may seem excessive, it was required to identify many of the weaker crosslinked RNA targets with confidence. Curiously, some of the strongest crosslinking peaks were located on transcripts of three RNase P RNA-like genes, *Rprl1-3*, which are homologous to but shorter than *Rpph1* (Figure 3B)^49, 50^. Although these transcripts have not been functionally characterized, they are expressed in a highly tissue-specific manner and are speculated to represent, like RNase MRP, a case of functional diversification of RNase P^49, 50^.

Our evidence for *in vivo* association of *Rprl1-3* transcripts with multiple subunits of RNase P but not the RNase MRP-specific Nepro suggests regulatory potential of the formed RNPs, including catalytic competence. This is noteworthy since homologs of *RPPH1* as well as *RMRP* have been found in other higher vertebrates, including humans^49, 50^.

Our study finds essentially all canonical as well as numerous novel candidate RNA substrates for each endonuclease, totaling to 80 for RNase MRP and 311 (including 124 tRNAs) for RNase P, and spanning a diversity of different RNA biotypes (Figure S3C). The identified target RNAs along with their putative cleavage sites provide a valuable resource for much further research into the enigmatic *in vivo* functions of these essential RNPs. Our findings suggest that both RNase MRP and RNase P have been exapted for a far wider variety of additional roles in mammalian cells than previously thought. We speculate that these newly acquired functions contribute to the pressure to preserve these enzymes and continue their evolution. It is also conceivable that the human pathologies associated with mutations in RNase MRP RNA and NEPRO are influenced by impairment of such functions^5, 6, 23, 51, 52^. Our characterization of the RNase MRP-specific subunits, definition of the RNA targeting repertoires of RNases P and MRP, and derivation of cell lines for inducible depletion of either enzyme provide valuable assets for future attempts to address these open questions.

## METHODS

### Cell culture and treatment

HEK293T cells were maintained in DMEM supplemented with 10% FBS (Cytiva, SH30910.03) and 100 U/ml Penicillin-Streptomycin (Gibco, 15140122) at 37°C and 5% CO2. mESC E14 cells were maintained in DMEM supplemented with 15% FBS (Cytiva, SH30910.03), 1% MEM NEAA (Gibco, 11140-050), 0.1% Beta-Mercaptoethanol (LifeTech, 21985-023), 10^6^ U/ml LIF (MiltenyiBiotec, 130-095-775) and 100 U/ml Penicillin-Streptomycin (Gibco, 15140122) at 37°C and 5% CO2.

To induce degron-mediated protein depletion, dTAG-13 (Tocris, 6605) and dTAG-V1 (Tocris, 6914) were dissolved in DMSO and used as an equimolar mixture at a final concentration of 500 nM (dTAG-13) and 1000 nM (dTAG-V1) for the indicated periods of time. For siRNA transfection, HEK293T cells were seeded in 6-well dishes and transfected 24 h later at about 40% confluence using the DharmaFECT 4 Transfection Reagents (Mirus, T-2001-01) with a pool of siRNAs targeting C18ORF21 (Horizon, M-016886-01-0005) or a non-targeting siRNA pool (Horizon, D-001206-13-05) at 50 nM.

To monitor the rate of cell growth, cells were seeded at 5,000 per well in a 48-well plate. At each indicated time point, cells were harvested by digestion with 0.25% trypsin-EDTA (Gibco, 2661762) from individual wells, and then counted using a hemocytometer (FisherScientific, 0267151B). Cell viability was determined via staining with Trypan Blue (Sigma,T8154-100ML).

To derive Nepro- or RPP21-degradable cell lines, degron tag (FKBPF36V)-containing knock-in cassettes were introduced homozygously into the last coding exon of the *Nepro* gene in mESC E14 cells or into the first coding exon of the RPP21 gene in HEK293T cells, as previously described^39^. Modified plasmids pCRIS-PITChv2-BSD-dTAG (Addgene, 91792), pCRIS-PITChv2-dTAG-BSD (Addgene, 91795), and pX333 (for simultaneous delivery of the PITCh and gene-targeting gRNAs; Addgene, 64073) were introduced into the cells via transfection. The HEK293T Nepro-degradable cell line was generated via infection with lentivirus generated from the pLEX_305-C-dTAG plasmid (Addgene, 91798) to drive dTAG-inducible expression of Nepro, and modified pX333 (for delivery of NEPRO gene-targeting gRNAs; Addgene, 64073) were introduced into the cells via transfection to knockout the endogenous NEPRO gene.

HEK293T Flag-HA-RPP25, mESC E14 Flag-HA-Rpp25, 3xFlag-RPP14, E14 3xFlag-RPP21 and E14 3xFlag-Nepro cell lines were derived by infection with lentivirus made from modified plasmids pUltra-hot (Addgene,24130) and then cell sorted based on expression of the Cherry protein. mESC E14 Flag-HA-Nepro and Flag-HA-C18orf21 cell lines were derived by infection with lenti-virus made from modified plasmids pLV-EF1a-IRES-Puro (Addgene 85132) and then puromycin selected.

### SDS-PAGE and western blot analysis

Western blots were performed as described previously^53^. Briefly, whole cell lysates were run on 15% SDS-polyacrylamide gels and transferred to supported nitrocellulose membrane (Bio-Rad) at 145 V for 1.5 h. Membranes were then blocked for 1 hour in 5% non-fat dry milk in TBST (137 mM NaCl, 20 mM Tris, pH 7.6, 0.1% Tween-20), rinsed three times for 5 min with TBST, and incubated with primary antibody (1:1000 in 3% BSA in TBST) overnight at 4 °C. Blots were washed in TBST 5 min for 3 times, incubated with HRP-conjugated secondary antibodies in 5% milk in TBST for 1 hour and then washed again before developing. Beta-Actin was detected by incubating blots with HRP-conjugated beta-actin antibody (1:15,000 in 5% milk in TBST) for 15 min followed by washing and developing. HRP signal was detected by developing with SuperSignal™ West Pico PLUS Chemiluminescent Substrate (Thermo Scientific, 34578).

### Immunofluorescence

Immunofluorescence was performed as previously described^53^. Briefly, cells were fixed in 4% formaldehyde in PBS for 10 min at room temperature, permeabilized in 0.5% Triton X-100 in PBS, blocked in 5% FBS in PBS, and probed with primary antibodies at 4°C for overnight. After an overnight incubation, the cells were probed with fluorochrome-conjugated secondary antibodies for 1 hour at room temperature and mounted using VECTASHIELD Antifade Mounting Medium with DAPI (Vector Laboratories, H-1200-10). Images were taken with the LEICA DMi8.

### Antibodies

NPM1 (Santa Cruz Biotechnology, sc-70392), RPP20 (Santa Cruz Biotechnology, sc-244043), RPP40 (Proteintech, 11736-1-AP), RPP21 (Proteintech, 16377-1-AP), RPP25 (Sigma, HPA046900-100UL), C18orf21 (Sigma, HPA065505-100UL), HA (Cell Signaling Technology, 3724S), FLAG M2 (Sigma, F1804-200UG),alpha Tubulin (Santa Cruz Biotechnology, sc-5286) and beta Actin (Sigma, A3854) were also used in this study.

### qPCR analysis

qPCR was performed as previously described^53^. Briefly, total RNA was extracted from samples using TRIzol Reagent (ThemoFisher, 15596018) and Direct-zol RNA Miniprep (Zymo Research, R2050) according to the manufacturer’s instructions. cDNA was prepared from equal amounts of RNA using PrimeScript RT Reagent Kit (Takara, RR037A) following manufacturer’s instructions. qPCR was performed using PowerUp SYBR Green Master Mix (ThermoFisher, A25742) to amplify the cDNA on the CFX Connect Real-Time PCR Detection System at the annealing temperature of 63°C. Relative mRNA levels were normalized to relative expression levels of the ACTIN gene that was used as an internal control.

#### Northern analysis

Northern blots with agarose/formaldehyde gels were performed as described previously^34^. All Northern probes were DNA oligonucleotides (ThermoFisher) 5’ end-labeled with ^32^P gamma-ATP (PerkinElmer, NEG502A250UC) and T4 polynucleotide kinase (New England Biolabs, M0201L) according to the manufacturer’s instructions. Briefly, for high-molecular-weight RNAs, 10–20 µg of total RNA was heated for 5 min at 70°C in 0.4 M formaldehyde/50% formamide/1× TT (30 mM tricine, 30 mM triethanolamine) /0.5 mM EDTA, cooled to room temperature in a thermocycler, and separated on 1% agarose/0.4 M formaldehyde /1× TT gels (15 cm × 15 cm × ∼0.5 cm) in 1× TT at 130 V for 5 min followed by 100 V for 3.5 h. The gels were rinsed with deionized water, soaked while gently shaking for 15 min in 50 mM NaOH, rinsed again with deionized water, and while gently shaking in 6× SSC for 15 min. RNA was capillary-transferred to Hybond N+ membranes (Cytiva, RPN203B) in 6× SSC overnight at room temperature. After transfer, membranes were cross-linked twice at 1200 mJ/cm2 before prehybridization in 10 mL of the hybridization buffer (Invitrogen, AM8670) for 60 min at 42°C. Probes (ITS1_3′end, mITS1_3′end) were then hybridized with the membrane overnight at 42°C. The next day, the membrane was washed sequentially with the wash buffer 1 (2xSSC, 0.1% SDS), wash buffer 2 (1xSSC, 0.1% SDS), and wash buffer 3 (0.1xSSC), once each for 30 min. Then membrane was developed using the storage phosphor screen (GE Healthcare, 0146-931) in an autoradiography cassette (Fisher Scientific, FBAC810) overnight at -20°C. Then next day, the phosphor screen was imaged with Typhoon imager(Cytiva).

### RNA fluorescence in situ hybridization

FISH probes were ordered from Integrated DNA Technologies. The HEK293T Nepro-degradable cells treated with DMSO or dTAG and HEK293T transfected with siCTRL or siC18ORF21 were hybridized with the RNA FISH Probe set labeled with CY5 or CY3, following the manufacturer’s instructions available online at www.biosearchtech.com/stellarisprotocols.

### Co-immunoprecipitation from cell lysates

The co-IP experiment was performed as previously described^53^. Briefly, mESC E14 cells were harvested, washed once with PBS, and lysed in whole-cell lysis buffer (20 mM HEPES-KOH pH 7.9, 10% glycerol, 300 mM KCl, 0.1% IGEPAL, 1 mM DTT) supplemented with cOmplete Protease Inhibitor Cocktail (Millipore Sigma, 11697498001) for 30 min at 4 °C. Supernatants were cleared off debris by a 30-min centrifugation at 17,000 × g at 4 °C. The lysates were then mixed with an equal volume of no-salt lysis buffer (20 mM HEPES-KOH pH 7.9, 10% glycerol, 0.1% IGEPAL, 1 mM DTT) supplemented with cOmplete Protease Inhibitor Cocktail to lower the final salt concentration to 150 mM KCl (IP buffer), added to ANTI-FLAG® M2 Affinity Gel(Millipore Sigma, A2220), and rotated for 2 h at 4 °C. After the IP, the beads were washed thoroughly with the IP buffer and the bound proteins were eluted with 150 μg/ml Flag (DYKDDDDK) peptide (GenScript, RP10586) in thermomixer at 4 °C, shaking at 1250 rpm for 1 h. The eluates were analyzed by western blotting.

### Protein complex purification

To purify protein complexes of Flag-HA RPP25, Flag-HA-Rpp25, 3xFlag-RPP14, E14 3xFlag-RPP21, E14 3xFlag-Nepro, Flag-HA-Nepro and Flag-HA-C18orf21, approximately 300 million HEK293T or E14 cells per experiment were harvested, flash-frozen in liquid nitrogen, and stored at −80 °C until use. Cells were resuspended in buffer A (20 mM HEPES-KOH pH 7.9, 10% glycerol, 300 mM KCl, 0.1% IGEPAL, 1 mM DTT, cOmplete Protease Inhibitor Cocktail; 100 μl of buffer A was used per 10^6^ cells) and rotated at 4 °C for 30 min. The lysates were centrifuged at 17,000 × g for 30 min at 4 °C, supernatants were collected, and then 250 μl ANTI-FLAG® M2 Affinity Gel (Millipore Sigma, A2220) was added and the mixture was rotated for 2 h at 4 °C. The affinity gel was then washed with 4 times buffer 1 (20 mM HEPES-KOH pH 7.9, 10% glycerol, 300 mM KCl, 0.1% Tween20, 1 mM DTT, cOmplete Protease Inhibitor Cocktail) and followed by 4 times buffer 2 (20 mM HEPES-KOH pH 7.9, 10% glycerol, 150 mM KCl, 0.1% Tween20, 1 mM DTT, cOmplete Protease Inhibitor Cocktail). The bound proteins were eluted with Flag peptide (150 μg/ml; GenScript, RP10586) in thermomixer at 4 °C, shaking at 1250 rpm for 1 h. The eluate of Flag pulldown proteins analyzed by mass spectrometry. For tandem affinity purification (TAP), the Flag eluate was mixed with anti-HA magnetic beads (Thermo Fisher Scientific, 88837) and rotated for 2 h at 4 °C. The beads were washed with 4 times buffer 2 and proteins were eluted using HA peptide (200 μg/ml; GenScript, RP11735) by shaking the beads in thermomixer at 1400 rpm for 45 min at 30 °C. The eluate was then analyzed by mass spectrometry.

### Immunoprecipitation mass spectrometry

Half of the elution sample was combined with 10mM DTT and 5% SDS, then incubated at 95 °C for 10 minutes to reduce and denature proteins. Following SP3 sample preparation^54^, SP3 beads and 100% acetonitrile were added to the samples to precipitate proteins. SP3 beads were washed twice separately with pure acetonitrile and 70% ethanol to clean up the sample. They were resuspended with 40µl 100mM triethyl ammonium bicarbonate (TEAB). The sample was digested overnight in 2µl trypsin (200 ng/µL) at 37 °C. The trypsin-digested sample was dried down to concentrate and resuspended in 4 µl 0.1 Formic Acid in H2O.

For LC-MS/MS analysis, 1µl of each sample was analyzed by the Exploris 480 mass spectrometer coupled with nLC Vanquish Neo UHPLC. Peptides were separated using a 25 cm × 75 µl column (Aurora Ultimate C18 1.7μm). The flow rate was 200 nl/min on 70-minute gradient (65 mins active gradient). The Solent A was 0.1 formic acid (FA) in water, the solvent B was 80% of acetonitrile with 20% 0.1 FA in H2O. The gradient steps are as follows: 1) 14 – 36% over 65 min for tryptic peptides separation; 2) 95% B for 5 mins to remove the residues from the column.

The MS/MS-based proteomics data was acquired on an Orbitrap Exploris 480 mass spectrometer operated in DDA mode. The ion source was operated in the positive ion mode with 1700 V spray voltage. The temperature of the ion transfer tube was 250 °C. The full scan parameters were as follows: MS1 scans with 120,000 resolutions with a scan range of 350–1200 m/z; auto mode of Maximum injection time; standard AGC target. Precursors were selected for fragmentation subject to intensity threshold of 5×10^3^, charge states between +2 and +5, and the exclusion duration was 45 seconds. The Data-Dependent MS2 scan was performed with a cycle time of 1 second. The MS2 scan was set to 0.7 m/z isolation window, 36% HCD collision energies, 45,000 resolution, auto mode of maximum injection time and 200% AGC target. The raw data were analyzed by MaxQuant (v2.6.1.0). The mouse protein database was downloaded from UniProt (2025). The maximum missed cleavages were set up to 2 for tryptic peptides. Oxidation (M) and acetylation (Protein N-term) were added as variable modification. The maximum number of modifications per peptide was set up to 5. The 5% peptide FDR and 5% Protein FDR were applied before summarizing peptides into proteins.

### RNA Immunoprecipitation and qPCR (RIP-qPCR)

Briefly, mESC WT, 3xFlag-RPP14, E14 3xFlag-RPP21, E14 3xFlag-Nepro, and Flag-HA-C18orf21 cells were seeded on 10cm plates, then harvest at 80% density with twice wash of PBS and pelleted by centrifugation at 1200 rpm for 5 minutes. Cell pellets were resuspended in 1000 µl of pre-chilled buffer A buffer A (20 mM HEPES-KOH pH 7.9, 10% glycerol, 300 mM KCl, 0.1% IGEPAL, 1 mM DTT, cOmplete Protease Inhibitor Cocktail) and rotated at 4 °C for 30 min. The lysates were centrifuged at 17,000 × g for 30 min at 4 °C, supernatants were collected and aliquot 50 µl of each sample were combined with 500 ml of TRI Reagent and stored at −80°C for subsequent RNA purification as input controls. Then 50 μl ANTI-FLAG® M2 Affinity Gel(Millipore Sigma, A2220) was added and the mixture was rotated for 2 h at 4 °C. The affinity gel was then washed with 4 times buffer 1 (20 mM HEPES-KOH pH 7.9, 10% glycerol, 300 mM KCl, 0.1% Tween20, 1 mM DTT, cOmplete Protease Inhibitor Cocktail) and followed by 4 times buffer 2(20 mM HEPES-KOH pH 7.9, 10% glycerol, 150 mM KCl, 0.1% Tween20, 1 mM DTT, cOmplete Protease Inhibitor Cocktail). The bound RNA and proteins were eluted with Flag peptide (150 μg/ml; GenScript, RP10586) in thermomixer at 4 °C, shaking at 1250 rpm for 1 h.For each sample, 50 µl of the Flag eluate were combined with 500 ml of TRIzol Reagent (ThemoFisher, 15596018) and stored at −80°C for subsequent RNA isolation. RNA was isolated Direct-zol RNA Miniprep (Zymo Research, R2050) according to the manufacturer’s instructions. cDNA was prepared from equal amounts of RNA using PrimeScript RT Reagent Kit (Takara, RR037A) following manufacturer’s instructions. qPCR was performed using PowerUp SYBR Green Master Mix (ThermoFisher, A25742) and the oligonucleotide primers to amplify the cDNA on the CFX Connect Real-Time PCR Detection System at the annealing temperature of 63°C. Relative mRNA levels were normalized to relative expression levels of the Actin gene that was used as an internal control. The relative enrichment of *Rpph1* and *Rmrp* in RNA purified from subunit-overexpressing cells was quantified by normalizing their levels to inputs and referencing them to the enrichment in non-overexpressing (WT) cells.

### AlphaFold prediction methods and Foldseek searches

Predictions of individual protein subunits of RNases P or MRP were generated with AlphaFold 2^55^ via the Google Colab notebook (https://colab.research.google.com/github/sokrypton/ColabFold/blob/main/AlphaFold2.ipynb)^56^ using the default parameters but specifying to relax all predicted structures using Amber. Predictions of the human RNase MRP were generated with AlphaFold3^57^ (https://alphafoldserver.com/welcome) using the default settings. Ten structures were predicted after providing the full-length sequences of the RNases P or MRP subunits, and aligned with ChimeraX^58^ to assess prediction convergence. ChimeraX was also used to prepare all structural figures.

We used the Foldseek web interface (https://search.foldseek.com/search), using as queries the experimentally determined structures of *S.cerevisiae* Rmp1 and Snm1 proteins (PDB: 7C7A)^8^.

### Individual-nucleotide resolution UV-crosslinking and immunoprecipitation (iCLIP)

All iCLIP experiments were performed in replicates following the iCLIP2 protocol^27^. Briefly, E14 mESCs stably expressing Flag-tagged Nepro, Rpp14, Pop5, or Rpp21 were grown in 10 cm plates and harvested at 85% confluence. The cells were washed with ice-cold PBS and irradiated with UV-light at 254 nm on ice. The irradiated cells were scraped, aliquoted into three 2-ml tubes and centrifuged at 5,000 g for 2 min at 4 °C. The supernatant was removed and the cell pellets were flash-frozen in liquid nitrogen and stored at -80 °C until use. Immunoprecipitation of the crosslinked RNP complexes was carried out using anti-Flag antibody (Millipore Sigma, F1804). The complete iCLIP experiment, including deep sequencing of the prepared cDNA libraries, was repeated in four replicates for each subunit.

### iCLIP data processing

The iCLIP data was processed essentially as previous described^59^. Briefly, FastQC (v0.11.9, https://www.bioinformatics.babraham.ac.uk/projects/fastqc/) was used to assess the data quality. High quality data were provided with FASTX Toolkit (v0.0.13, http://hannonlab.cshl.edu/fastx_toolkit/. parameters: -Q 33 -q 10 -p 100). Flexbar (v3.5.0)^60, 61^ was utilized to demultiplex data based on their indexes, followed by mapping to the mm10 genome with STAR genome aligner v2.7.3a^62^. Umi_tools v1.0.1^63^ was used to remove PCR duplications. High confident cross link sites were identified with PureCLIP v1.3.1 (parameter: - ld -nt 8)^64^. The crosslink sites identified in at least two replicates were kept and expanded to a ∼9-nt region as described previously^59^. Binding sites were annotated using GENCODE (vM25) comprehensive gene annotation. Bigwig files were produced with bedtools v2.31.1^65^ by shifting the reads 1 nucleotide upstream to identify the genuine crosslink sites and by normalizing to the whole library with RPM (reads per million). The IGV version 2.9.4^66, 67^ traces shown in the figures were summaries of all four replicates. RNase MRP specific peaks were those identified in Nepro, Rpp14, Pop5, but not in Rpp21 iCLIP datasets, whereas RNase P specific peaks were those identified in Rpp21, Rpp14, Pop5, but not Nepro iCLIP datasets.

## ACKNOWLEDGEMENTS

We thank members of the Murn and Cheloufi laboratories for helpful discussions. We are grateful to Mary Hamer, David Carter, and Melanie Oakes for assistance with experiments. This work was supported in part by a grant from the US National Science Foundation (2054195) to J.M. and S.C. and a MIRA award from the NIGMS of the NIH (R35GM148218) to N.S. S.H. and M.C. were supported by the UCR Stem Cell Center TRANSCEND training CIRM grants. J.M. is supported by the U.S. National Institutes of Health grant GM144693 and S.C. is supported by the U.S. National Institutes of Health grant GM151004.

## AUTHOR CONTRIBUTIONS

J.M. conceptualized the study. Y.L., S.H., and J.M. designed experiments and data analyses. Y.L., S.H., K.P., S.Z., M.C., and J.M. performed all experiments. S.H. performed bioinformatic analyses. S.C., N.S., and W.F.M. contributed key reagents and provided guidance. J.M. acquired funding and supervised this work. J.M. wrote the manuscript with input from all other authors.

## Supplemental Figures

**Supplemental Figure 1, related to Figure 1.**
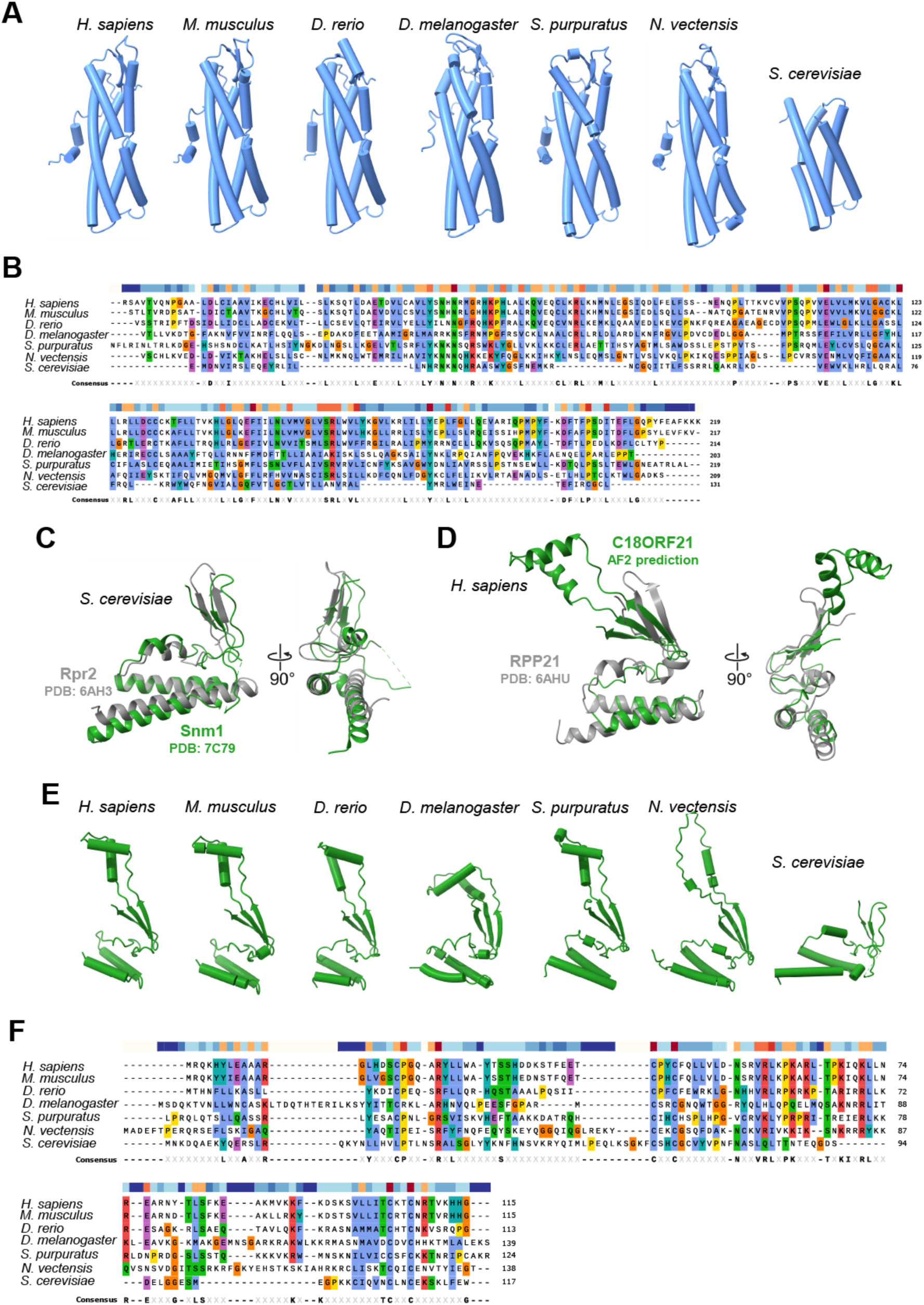
Structural homology of known and predicted subunits of RNases P and MRP. **(A)** A representative AF2 prediction of the N-terminal helical bundle of NEPRO for each of the shown NEPRO orthologs (*Homo sapiens*, human; *Mus musculus*, house mouse; *Danio rerio*, zebrafish; *Drosophila melanogaster*, fruit fly; *Stronglycentrotus purpuratus*, sea urchin; *Nematostella vectensis*, starlet sea anemone; *Saccharomyces cerevisiae, brewer’s yeast*). **(B)** Sequence alignment of the N-terminal helical bundle of NEPRO orthologs shown in (A) (performed with SnapGene, version 4.3.11). A conservation score is indicated above the alignment. Blue, yellow, and red colors indicate low, median, and high conservation, respectively. A consensus sequence is provided below the alignment. **(C)** Superposition of the N-terminal helices and beta-sheets of the yeast Rpr2 (PDB: 6AH3) and Snm1 (PDB: 7C79). **(D)** Superposition of the N-terminal helices and beta-sheets of the human RPP21 (PDB: 6AHU) and C18ORF21 (AF2 prediction). **(E)** A representative AF2 prediction of the N-terminal helices and beta-sheets of C18ORF21 for each of the shown C18ORF21 orthologs, as in (A). **(F)** Sequence alignment of the N-terminal helices and beta-sheets of C18ORF21 orthologs shown in (E). See also (B).

**Supplemental Figure 2, related to Figure 1.**
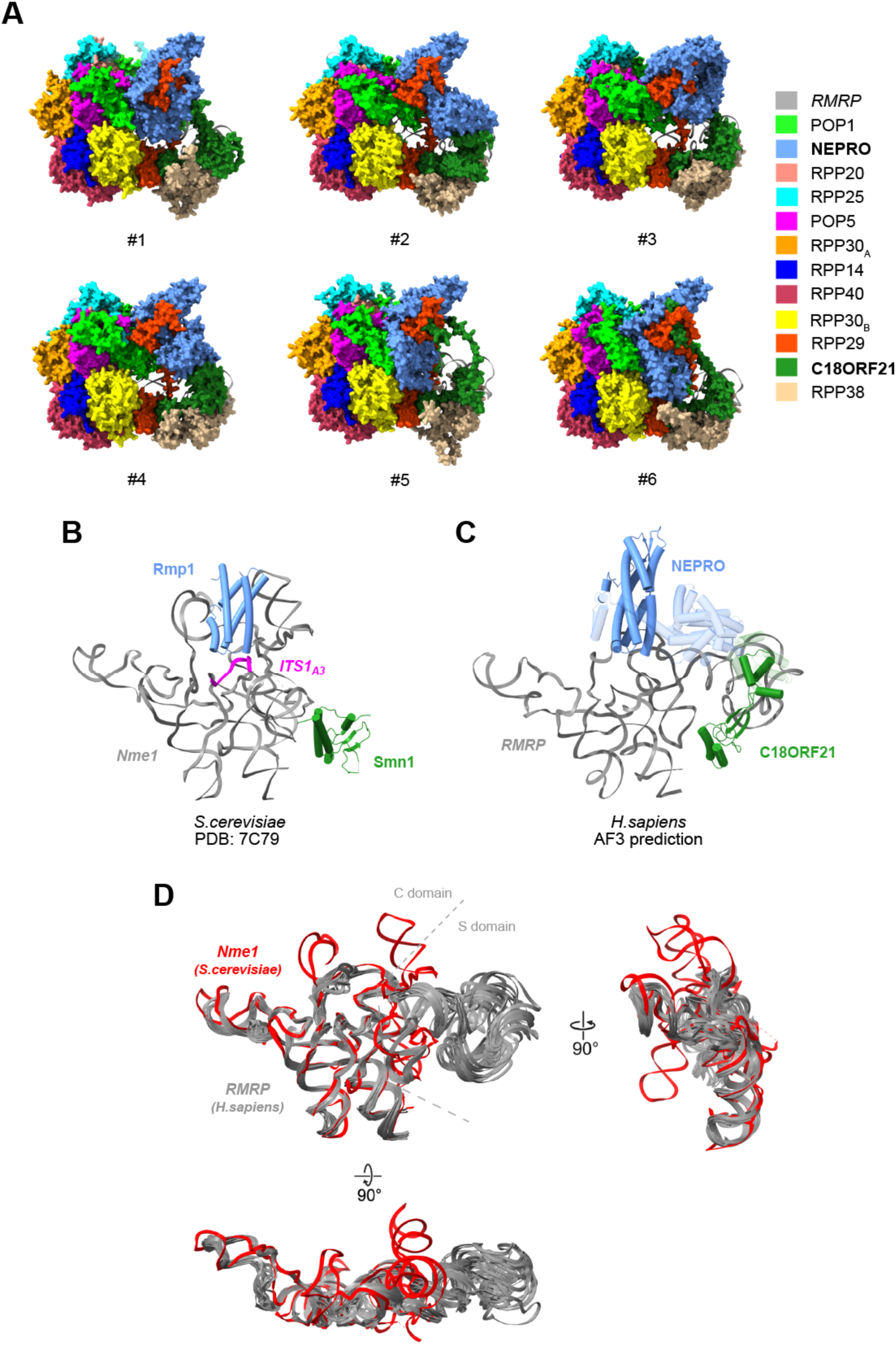
Structural predictions of the human RNase MRP RNP. **(A)** Six AlphaFold3 (AF3) predictions of the human RNase MRP complex using full-length sequences of the listed subunits. Our identified protein subunits, NEPRO and C18ORF21, are highlighted in bold. **(B)** Cryo-EM structure of the yeast RNase MRP in complex with its RNA substrate, ITS1_A3_ (PRB: 7C79). Rmp1, Snm1, *Nme1*, and *ITS1_A3_* are shown as visible in the structure. All other subunits are hidden. **(C)** AF3 prediction #4 in (A), showing only *RMRP* and full-length NEPRO and C18ORF21. The converging protein segments from 10 independent predictions are shown opaque and the variable segments are transparent. **(D)** Ten AF3 predictions of the human RNase MRP and the yeast cryo-EM structure of RNase MRP (PDB: 7C79; red) were aligned on *RMRP*. Shown are only *RMRP* and *Nme1*. The gray dashed lines denote the approximate boundary between the C and S domains in *RMRP*.

**Supplemental Figure 3, related to Figure 3.**
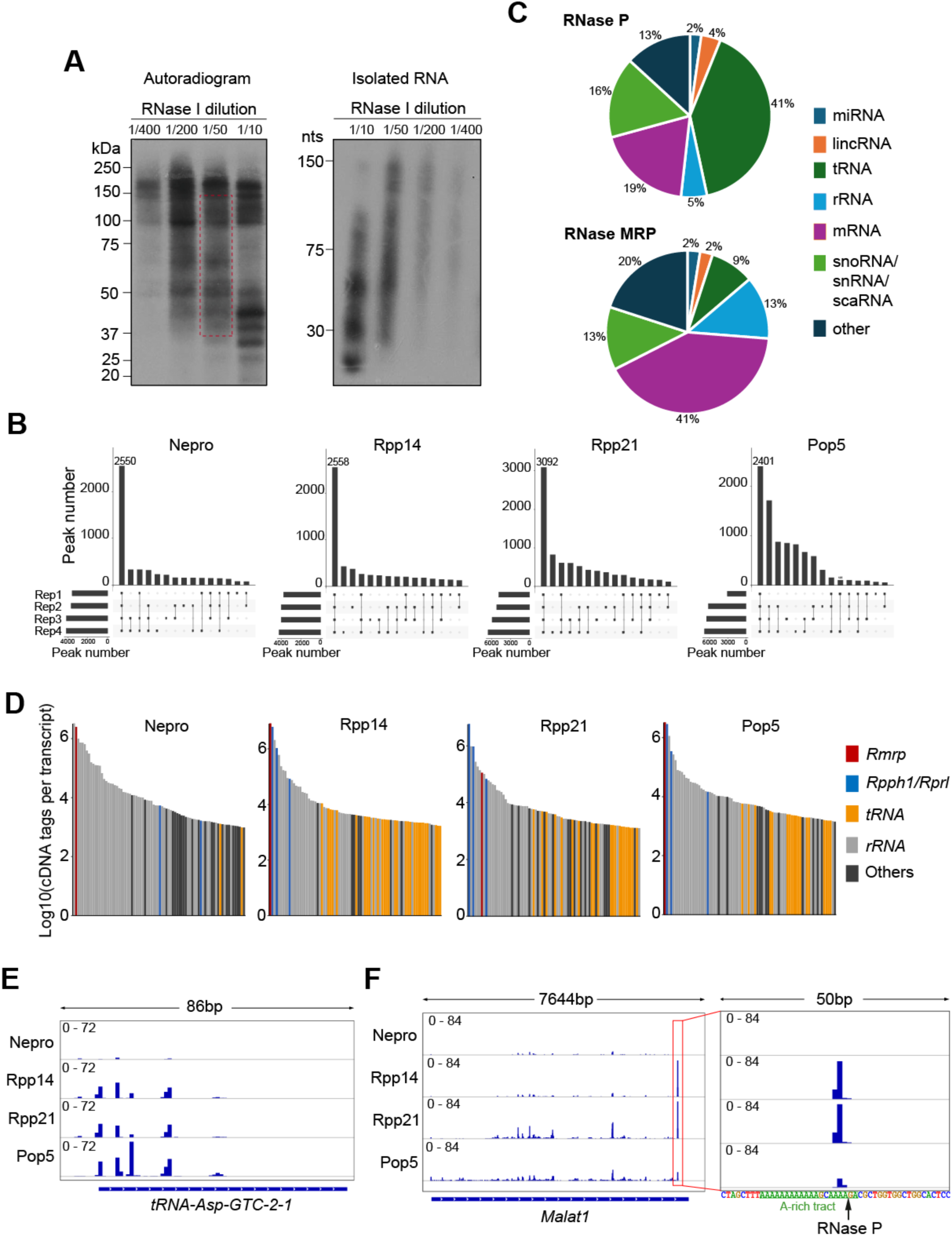
RNA-binding specificity of RNases P and MRP. **(A)** Crosslinking of RNases P and MRP to RNA in mESCs. (*Left*) Autoradiogram of crosslinked and labeled protein/RNA complexes precipitated via Flag-tagged Rpp14. The experiment was intended for optimization of RNase I dilution. A similar pattern of radioactive bands was seen with iCLIPs of other subunits. Dashed red rectangle indicates the approximate range of crosslinked RNP complexes that were cut from the membrane and used for preparation of iCLIP libraries. (*Right*) Radioactively labeled RNA extracted from the crosslinked complexes on the left. **(B)** Reproducibility of the binding site (peak) calling in different iCLIP datasets. Four replicates were performed for each subunit-iCLIP experiment, and the binding sites were identified separately. Horizontal bars indicate peak numbers identified in each replicate experiment. Bar plots and dot plots indicate peak number called in different sets of replicates. **(C)** Biotype distributions of the 311 RNase P-bound and 80 RNase MRP-bound transcripts. **(D)** Eighty most highly bound transcripts in each iCLIP dataset. iCLIP cDNA tags within all binding sites on a transcript were summarized and ordered from highest (left to lowest (right) in each plot. Gencode vM25 annotations of individual transcripts are shown. **(E)** Example of a crosslinking profile on a tRNA transcript (*tRNA-Asp-GTC-2-1*). **(F)** Crosslinking profile on *Malat1* shown for each iCLIP dataset. The highlighted binding peak in the left window is shown magnified on the right. Note the proximity of the peak to the annotated RNase P cleavage site within the A-rich tract (highlighted in green).

**Supplemental Figure 4, related to Figure 4.**
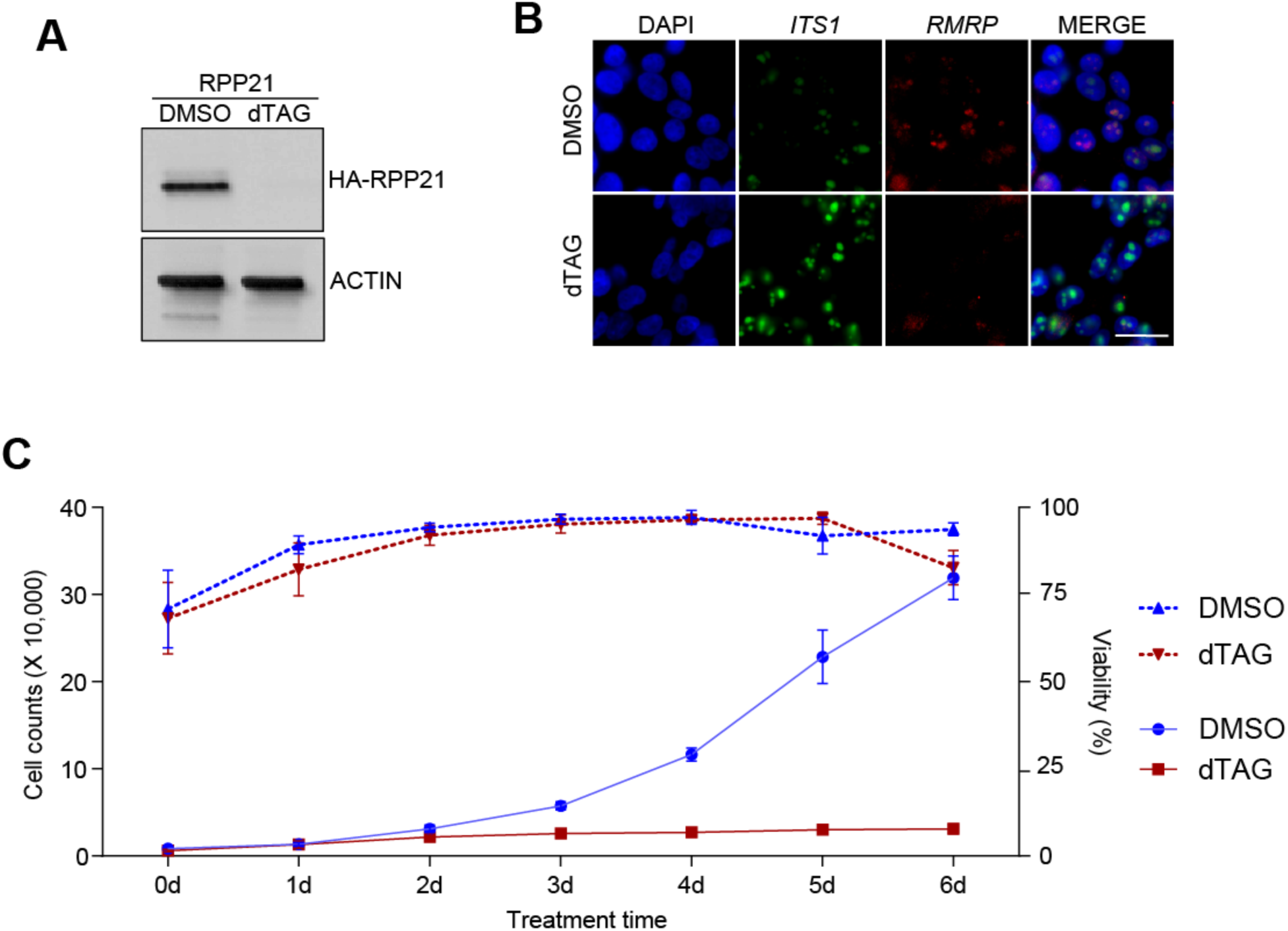
Requirement of NEPRO for pre-rRNA processing and cell proliferation. **(A)** Depletion of the endogenous, HA-tagged RPP21 in our engineered HEK293T cells at 2 days of treatment with dTAG (n = 3). **(B)** RNA FISH of *ITS1* and *RMRP* at 2 days of treating Nepro-degradable HEK293T cells with DMSO or dTAG. Scale bar, 25 µm. **(C)** Growth of Nepro-degradable HEK293T cells over the indicated periods of treatment either continuously with DMSO (blue; Ctrl) or with dTAG (red). Dashed lines indicate cell viability. Data are shown as mean ± SD (n >= 3).

